# Crispr/Cas9 Targeted Capture of Mammalian Genomic Regions for Characterization by NGS

**DOI:** 10.1101/372987

**Authors:** Alexei Slesarev, Lakshmi Viswanathan, Yitao Tang, Trissa Borgschulte, Katherine Achtien, David Razafsky, David Onions, Audrey Chang, Colette Cote

## Abstract

The robust detection of structural variants in mammalian genomes remains a challenge. It is particularly difficult in the case of genetically unstable Chinese hamster ovary (CHO) cell lines with only draft genome assemblies available. We explore the potential of the CRISPR/Cas9 system for the targeted capture of genomic loci containing integrated vectors in CHO-K1-based cell lines, and compare it to popular target-enrichment methods and to whole genome sequencing (WGS). The CRISPR/Cas9-based techniques allow for amplification-free capture of genomic regions, which reduces the possibility of sequencing artifacts. Other advantages of these methods are the ease of bioinformatics analysis, potential for multiplexing, and the production of longer sequencing templates for real-time sequencing. The utility of these protocols has been proven by identification of transgene integration sites and flanking sequences in a number of CHO cell lines. However, data produced by these and other targeted capture methods are not always sufficient to analyze complex genomic rearrangements (CGRs) or unexpected sequences introduced into genome by vector integration events. In contrast, WGS provides complete information about vector integration sites, vector copy number, CGRs, and foreign DNA-but despite these benefits, WGS is not easily implemented due to the cost and complexity of the analysis.

## INTRODUCTION

Considerable effort has been devoted to develop ‘target-enrichment’ methods, in which genomic regions are selectively captured from a DNA sample before sequencing. The rationale for those efforts is that sequencing the retained genomic regions is more time and cost-effective than whole-genome sequencing (WGS), with the added benefit of data being considerably less cumbersome to analyze.

The most common techniques that have existed for some time can be categorized according to the nature of their core reaction principle. In the “hybrid capture” method, nucleic acid strands derived from the input sample are hybridized specifically to prepared DNA fragments complementary to the targeted regions of interest, either in solution or on a solid support, so that one can physically capture and isolate the sequences of interest. There are several popular commercial solutions for this – SureSelect and HaloPlex from Agilent Technologies, xGen from Inegrated DNA Technologies (IDT), Ion TargetSeq from Life Technologies, – all adapted from the hybrid selection method originally developed by Gnirke *et al*^1^. These assays place a premium on performance and are cost effective if used for parallel characterization of the same genomic region in multiple samples. However, these methods become cost-prohibitive for one-time use on a single genomic locus. Also, the fragment sizes utilized by current DNA target enrichment technologies remain a limiting factor, producing DNA fragments generally shorter than 1 Kb. In the “selective circularization” methods (*e.g.* MIPs, gap-fill padlock probes, selector probes) single-stranded DNA circles containing the target region sequences are formed (by gap-filling and ligation chemistries) in a highly specific manner, creating structures with common DNA elements that are then used for selective amplification of the targeted regions. PCR amplification methods can be directed toward the targeted regions by conducting either multiple long-range PCRs in parallel, a limited number of standard multiplex PCRs, or using highly multiplexed PCR methods that amplify very large numbers of short fragments. However, these methods heavily rely on PCR amplification, which may result in artifact sequence junctions that are difficult to distinguish from real ones. Recently, a targeted locus amplification (TLA) method has been developed on the basis of the crosslinking of physically proximal sequences^2^. This technique allows re-sequencing and haplotyping of long multikilobase genomic regions, but the method is not easily implementable. Given the operational characteristics of these different targeted enrichment methods, they naturally vary in their suitability for different fields of application (reviewed in^3–5^).

The advent of engineered DNA-binding molecules, such as zinc finger proteins (including ZFNs), transcription activator-like effector (TAL or TALE) proteins^6^, and the clustered regularly interspaced short palindromic repeats (CRISPR) system^7^, has enabled high efficiency genome editing and *in vivo* chromatin immunoprecipitation (ChIP) techniques. In addition to the conventional *in vivo* ChIP, Fujita and Fujii reported that using enChIP with recombinant engineered DNA-binding molecules, such as a TAL protein, it is feasible to isolate a genomic region of interest from a cell^8^. Among those aforementioned engineered DNA-binding molecules, the CRISPR/Cas9 system is the most convenient, economical and time-efficient, and has rapidly risen to prevail over technologies with similar functions^9,10^. CRISPR/Cas9 has been used recently for targeted sequencing microsatellite-spanning sequences^11^ and loci associated with repeat expansion disorders in conjunction with Pacific Biosystems’ Single Molecule Real Time (SMRT) sequencing^12^.

In this study, we developed and tested several *in vitro* CRISPR/Cas9 protocols for targeted capture of transgene containing regions in genomic DNA (gDNA) from recombinant Chinese hamster ovary (CHO) cell lines. These protocols are target-specific and can be used for capturing any genomic region. Genomic instability is a hallmark of CHO cell lines^13–16^, thus necessitating constant monitoring of recombinant CHO clones following the random genomic integration of transgenes. We demonstrate that CRISPR RNA-guided endonucleases (RGENs) can be employed to enrich desired long DNA fragments from mixtures of purified genomic DNA, and then used for integration locus identification by next generation sequencing (NGS). We also compare targeted enrichment methods for locating vector integration sites and associated chromosome-wide rearrangements in recombinant CHO cell lines with whole genome sequencing (WGS) and discuss the best application of these methods.

## RESULTS

### CRISPR/Cas9 target capture protocols

*In vitro* CRISPR/Cas9 methods use biotin-labeled RNA-guided engineered nucleases (RGEN) to specifically bind to targeted regions and enable either the isolation of the targeted DNA fragments from the population of gDNA, or cleave the targeted regions and produce double-stranded breaks that can be further enzymatically labeled by biotin and isolated from the mixture of genomic fragments. The purified target DNA fragments are then used for NGS library construction and sequencing. Three different Cas9 RGEN protocols have been developed and tested using three CHO-K1-based cell lines, AE54SL, AD49ZG and AD49ZH, all bearing integrated vectors with immunoglobulin transgenes.

In both *RGEN-R* and *RGEN-TdT* protocols (Figure 1) gDNA is first sheared using routine methods to make long, > 20 Kb fragments, which are then treated with a DNA polymerase and ddNTP to block free 3’ DNA ends (*step 1*). Next, Cas9 nuclease is incubated with an appropriate single-guide RNA (sgRNA) to form RGEN that is then used to cleave ddNTP-treated DNA fragments at a targeted site, resulting in blunt-ended DNA fragments (*step 2*). In the *RGEN-R* protocol, a short double-stranded DNA (dsDNA) adapter with a site for a rare-cutter restriction enzyme and containing a biotin molecule at the free 5’-end is ligated to the cleaved DNA fragment (*step 3*). DNA fragments containing a ligated biotinylated adapter are then isolated from the DNA mixture using streptavidin-coated magnetic beads and released from the beads by digestion with the desired restriction enzyme (*step 4*). In the *RGEN-TdT* protocol, in place of the adapter ligation step, the DNA fragments are labeled by TdT-mediated dUTP-biotin end-labeling (TUNEL)^17^ and purified using streptavidin-coated magnetic beads. Single stranded DNA (ssDNA) fragments are released after heat or chemical treatment of the biotin-streptavidin complexes. Depending on the yield, in both protocols the released DNA fragments may require whole genome amplification using available chemistries before proceeding with NGS library construction (*step 5*).

**Figure 1.**
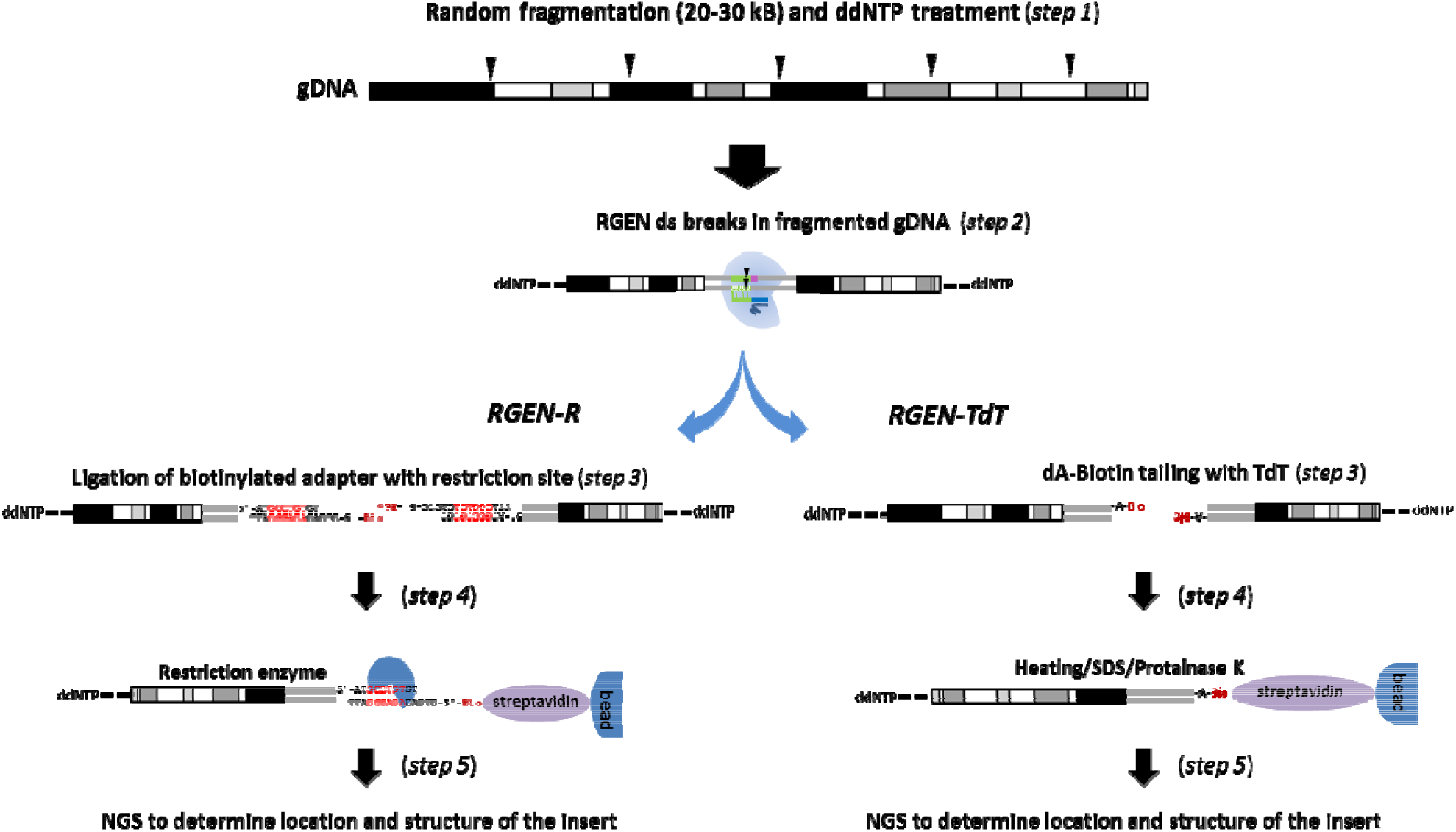
Schematic representation of main steps in the RGEN-R and RGEN-TdT protocols.

It is interesting to note that an earlier variant of the RGEN-TdT without a ddNTP pre-treatment step failed to meaningfully enrich the targeted loci. Only after introducing this step the RGEN-TdT protocol resulted in drastic enrichment of the vector integration region. It is apparent that pre-treatment of the genomic DNA with Klenow fragment and ddNTP prevents nonspecific biotin labeling of the DNA by TdT. Another useful step that increased the RGEN-TdT sensitivity is addition of dATP alongside with biotin-16-dUTP to the labeling mix to increase the number of biotin-modified nucleotides at the 3-OH DNA termini^17^. TdT prefers dATP as a substrate and can add up to five times more dATP than other nucleotides at an identical concentration. Hence, dATP was included in the labeling mix to increase the length of the end-labeling and consequently the number of biotin-modified nucleotides added to the termini of each DNA break. This increased the efficiency of DNA binding to streptavidin magnetic beads.

In the RGEN-D protocol 3’-biotinylated sgRNAs and a mutant, inactive Cas9 protein are used to generate biotin-labeled RGENs that can only bind, but not cleave targeted loci at the binding sites (Figure 2, *A*). Alternatively, a biotinylated, inactive Cas9 protein can also be used (Figure 2, *B*). Biotin-labeled RGENs (about one RGEN per 2-3 Kb stretch of DNA) are incubated with fragmented gDNA (~ 5 Kb fragments) and then RGEN-DNA complexes are isolated from the DNA mixture either using streptavidin-coated magnetic, followed by RNAse A and Proteinase K treatment to release dsDNA fragments. Purified DNA fragments are then used for TruSeq® Illumina® library construction and sequencing.

**Figure 2.**
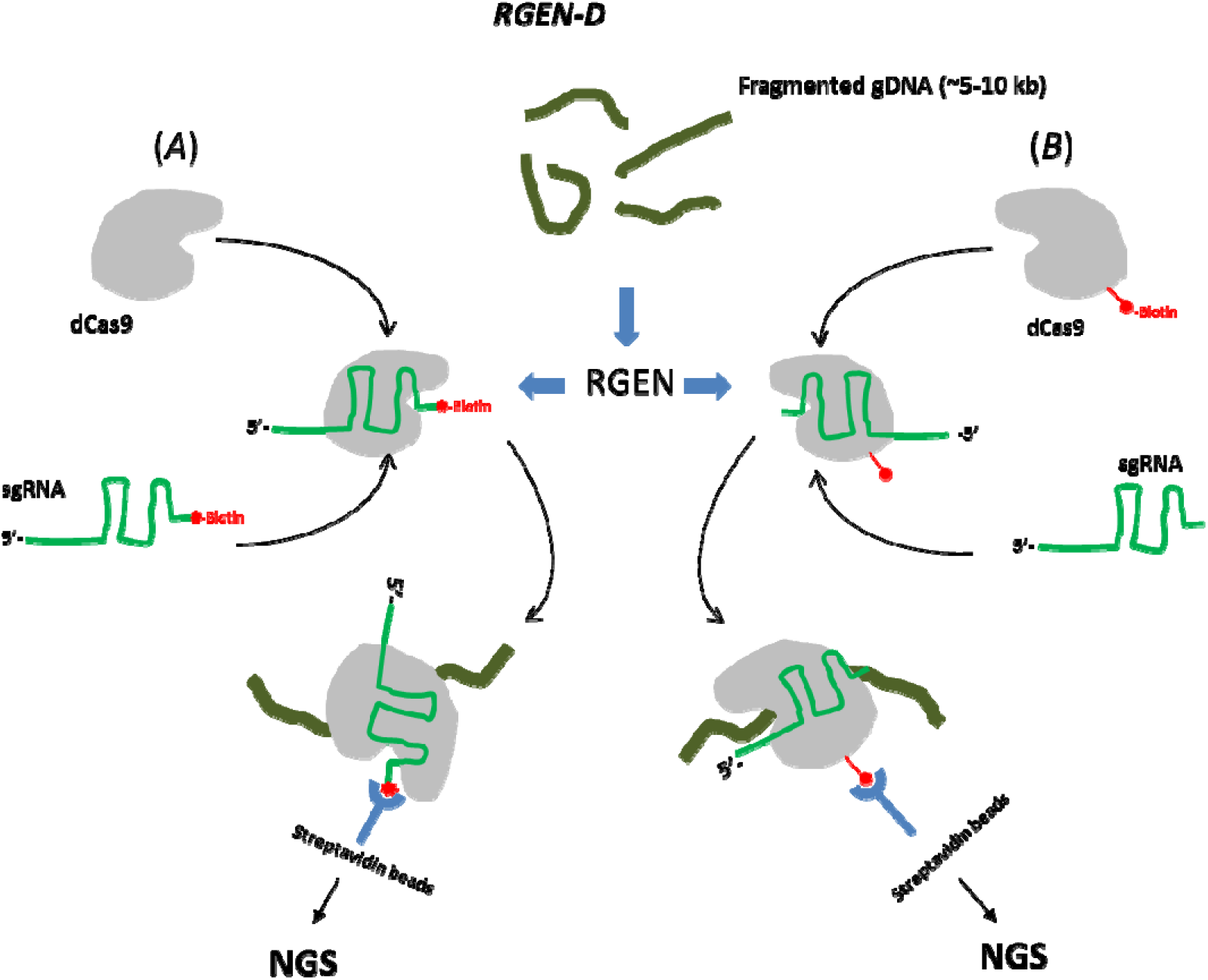
Schematic representation of main steps in the RGEN-D protocol.

A summary of several target enrichment experiments with three different protocols and using two different CHOK1-derived clones are presented in Table 1. All experiments were designed as proof-of-principal to demonstrate the utility of Cas9 proteins for selective enrichment of DNA. For the RGEN-R and RGEN-TdT protocols, a single target site for Cas9 cleavage was identified in the middle of the 11,489 bp integration vector (gRNA1), while five target sites were chosen for the RGEN-D protocol (Supplementary Table 1, Figure 4). To assess the enrichment efficiency, we used, in all cases, a comparative ∆∆C_T_ TaqMan Assay^18^. This assay was designed for estimating the copy number fold change of a targeted sequence in the enriched DNA sample relative to the target’s “normal” copy number in the control DNA sample.

**Table 1:** Fold change enrichment of targeted locus after RGEN treatment, calculated by ΔΔC_T_ method.

| Sample | AD49ZG |  |  |  | AD49ZH |  |  |  |
| --- | --- | --- | --- | --- | --- | --- | --- | --- |
| Protocol | RGEN-TdT |  | RGEN-D |  | RGEN-R |  | RGEN-TdT |  |
| Sample treatment | Control | Enriched | Control | Enriched | Control | Enriched | Control | Enriched |
| Cgr Spag5 average $C_T$ | 37.2±0.15 | ND | 24.5±0.15 | 29.8±0.16 | 25.21±0.15 | 35.85±0.19 | 30.1±0.12 | ND |
| Hsa IGHM average $C_T$ | 36.1±0.17 | 37.5±0.17 | 25.0±0.10 | 25.2±0.09 | 17.55±0.08 | 25.05±0.10 | 30.0±0.10 | 37.2±0.15 |
| $\Delta C_T$ Spag5-IGHM | 1.1±0.22 | ND | -0.5±0.18 | 4.6±0.18 | 7.66±0.17 | 10.80±0.21 | 0.1±0.16 | ND |
| $\Delta\Delta C_T$ ($\Delta C_T$ enriched - $\Delta C_T$ Control) | 0.0±0.22 | ND | 0.00±0.18 | 4.1±0.18 | 0.00±0.17 | 3.14±0.21 | 0.0±0.16 | ND |
| Fold difference in IGHM relative to Control | 1 (0.6-1.2) | 56.8* | 1 (0.6-1.1) | 17.1 (19.4-15.1) | 1 (0.6-1.1) | 8.8 (7.6-10.2) | 1 (0.6-1.1) | 29.2* |

\*Determined by NGS

Following isolation of the target DNAs by the described protocols, we used Illumina® sequencing by synthesis to characterize the targeted DNA regions and estimate the level of enrichment. The RGEN-TdT’s (AD49ZG and AD49ZH clones) and RGEN-D’s (AD49ZG clone) captured DNAs were sequenced using Illumina TruSeq Nano libraries on an Illumina® MiSeq instrument. Quality control and general statistics reports on the sequencing runs are available in the Supplementary Table 2. In the RGEN-D/AD49ZG sample the unenriched chromosomal regions show an average coverage of 0.5X, which can be considered the diploid baseline for the coverage comparisons. Thus, the integrated vector mean coverage of 16.9X in this case would correspond to ~17-fold enrichment of the vector region, assuming there are four vector copies and all of them are integrated into one chromosome of the chromosome pair (Table 2). This is close to the 17.1X value obtained by ∆∆C_T_ TaqMan method (Table 1). As seen in Tables 1 and 2, the highest enrichment between different RGEN protocols is consistently achieved using the RGEN-TdT protocol.

**Table 2.**
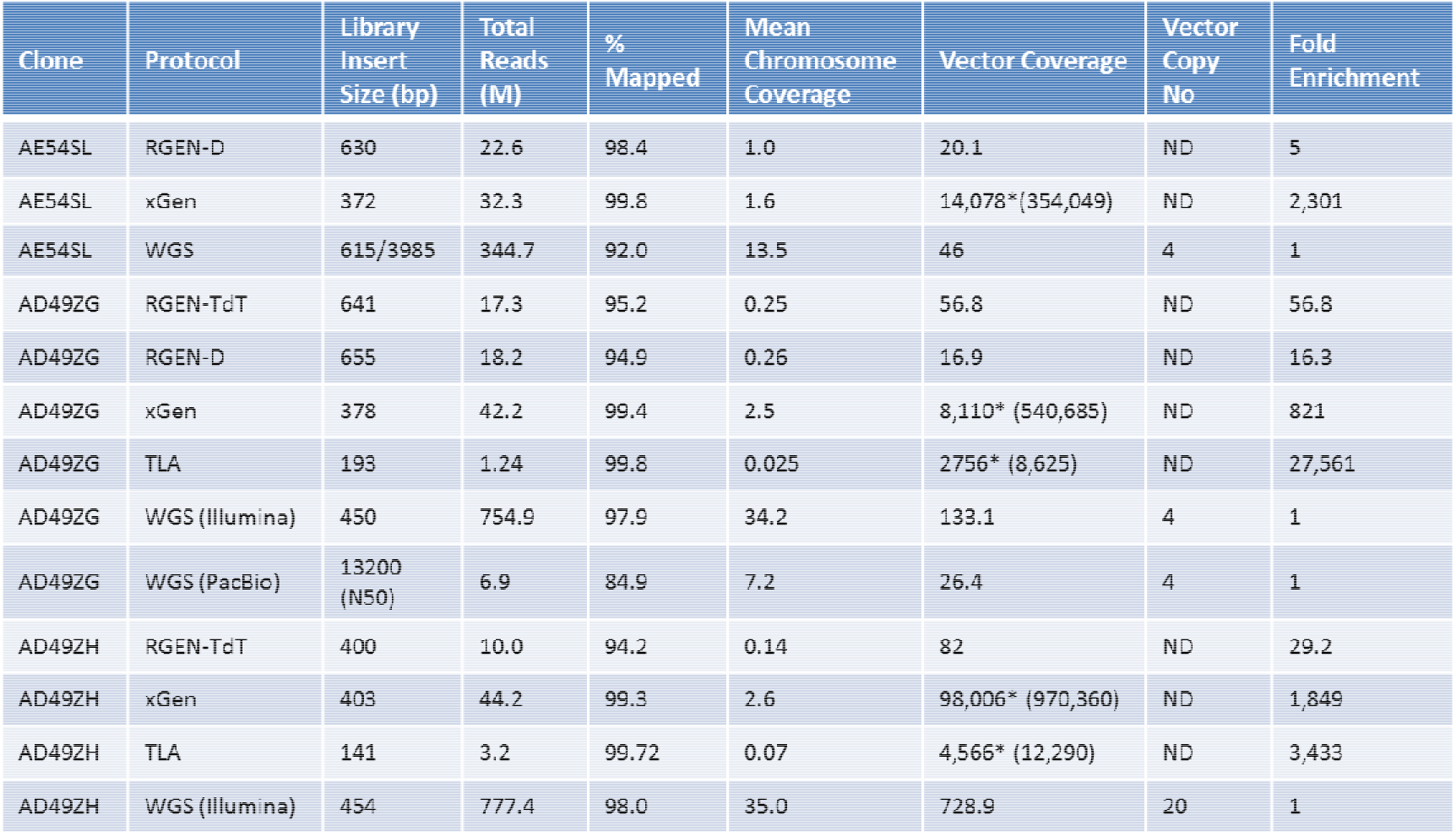
Summary of the NGS analysis of targeted capture experiments performed in this study. *Vector coverage after removal of PCR duplicates. PCR duplication rates were calculated according to Bansal^24^. The integrated vector copy numbers were calculated using the WGS Illumina® and PacBio data, as ratios of the vector coverage to the mean chromosome coverage. The fold enrichments were normalized by the corresponding vector copy numbers.

### Structure of the Integration Loci

Because RGEN protocols isolate all DNA fragments in close proximity to the vector integration site, it is possible to identify surrounding host system (*e.g.* CHO-K1) sequences, in addition to the adjacent vector sequence. The only challenge is the selection of proper reference genomes for mapping analysis. In the case of Chinese hamster and CHO cell lines it is difficult due to the lack of a complete reference genome assembly. Therefore, to characterize the sites of vector integration, we created custom reference genomes comprised of CHO-K1 chromosomes, a corresponding integration vector and *Escherichia coli* K-12 MG1655 (*E. coli*) fasta sequences as pseudochromosomes (see METHODS for details). The use of custom chromosome-based reference genomes has simplified the analysis and presentation of vector integration events, and allowed visualization of associated chromosomal rearrangements. Moreover, such an approach is fully compatible with popular structural variation (SV) callers and pipelines (SpeedSeq^19^, Manta^20^, Lumpy^21^, Delly^22^), allowing their usage without modification, obtaining results in the standard VCF format. We also used an in-house custom genomic stability (GS) pipeline to identify read pairs where one read is fully or partially mapped to a vector sequence, but paired or mate paired reads are unmapped. Assembling these reads followed by Basic Local Alignment Search Tool (BLAST^23^) analysis of resulting contigs against a reference genome results in identifying all vector|vector and/or vector|chromosome breakpoints. An example of this pipeline run is provided in the Supplemental Information.

To investigate the relevant performance of different capture protocols and the WGS method, we sequenced captured DNA from several CHO-K1-derived cell lines AE54SL, AD49ZG, and AD49ZH on an Illumina® MiSeq instrument (or using an Illumina® HiSeq2500 instrument, in case of WGS) (Table 2).

#### AE54SL cell line

Both the RGEN-D and xGen targeted capture protocols, as well as WGS, identified two vector fusion breakpoints in the AE54SL cell line (Figure 3). However, the vector copy number per genome can be estimated only from the WGS data (Table 2), based on the average aligned read depth of the Illumina® WGS datasets across the genomes (Supplementary Table 2), and which we consider the diploid baseline. Thus, a mean per chromosome coverage is ½ of the diploid baseline and this value is used for the vector copy number calculation in all cases (Table 2).

**Figure 3.**
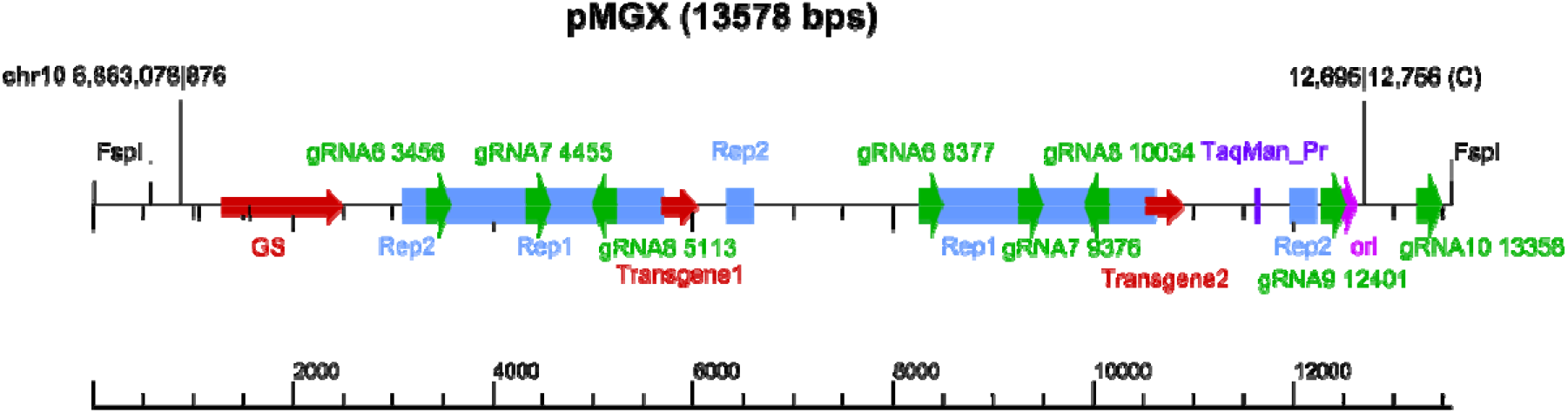
Map of the integration vector in the AE54SL cell line. Different vector features are shown color-coded. Sizes of guid RNAs used in the RGEN-D protocol are not to scale. Two identified fusion breakpoints are shown (chr10|vector and vector|vector junction (head-to-head)).

The xGen protocol (IDT) results in extremely high coverage across the vector, which is not surprising as this protocol relies on two rounds of NGS library amplification by PCR. It has therefore a great potential for multiplexing and/or can be run on a low output sequencer (*e.g*., on an Illumina® iSeq 100). However, it often results in PCR chimeras, which are impossible to separate from real fusion breakpoints without verification of all junctions by separate method(s). For example, it resulted in three false positives in this clone: fusions between positions 7276|6304C, and 7247|11081C (vector chimeras), as well as a fusion between 4152|6,881,753 (vector|CHO chr10 chimera). These specific chimeric junctions are absent in both the PCR-free RGEN-D and WGS data sets.

The only vector “head-to-head” junction together with the only chromosome-vector fusion breakpoint in this clone (Figures 3, 4) suggest a recombination event involving the vector (pGMX) in a head-to-head duplication with the upstream region of chromosome 10 (the letter of unknown size due to limitations in the technology). Using the corresponding WGS data, we looked more closely at this region, leveraging both depth of coverage (DOC) of chromosome 10 and discordant pair-end mapping techniques to establish a sequence of events that best explains the observed copy number and SVs.

**Figure 4.**
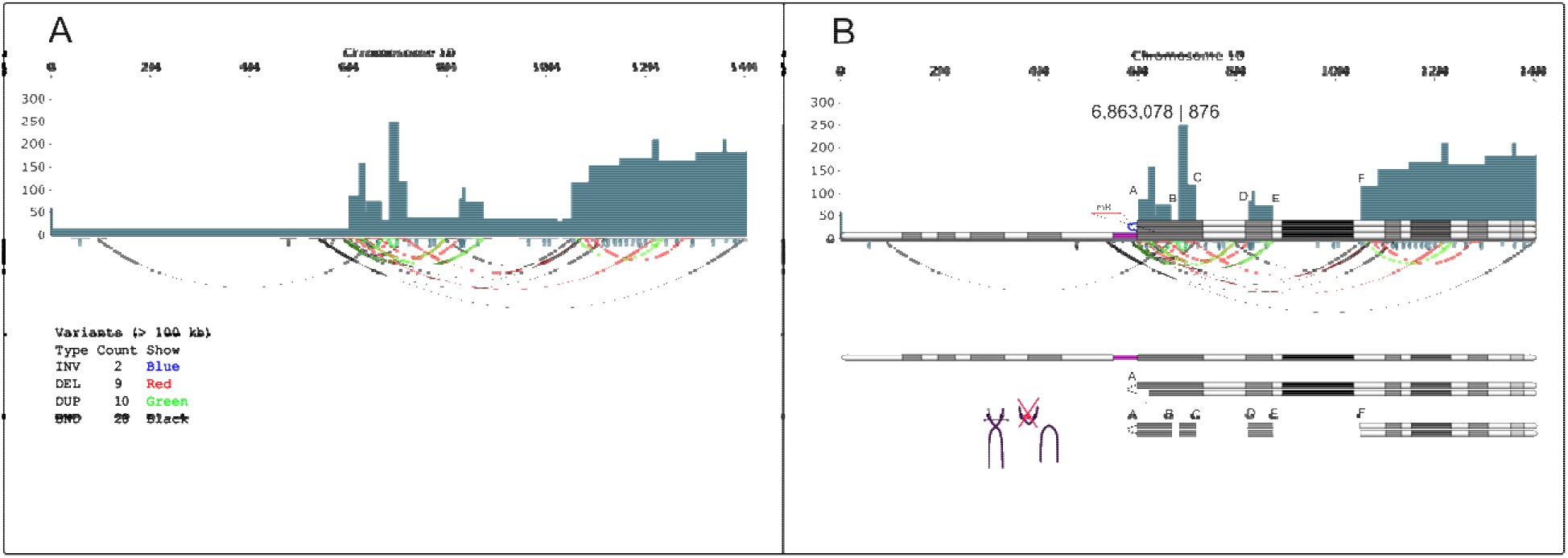
Reconstruction of the integration site rearrangements in the chromosome 10 of the AE54SL cell line. ***A,*** Copy number and structural variations for the chromosome 10 visualized by SplitThreader. Copy number segmentation is based on coverage averages in 10 kbp bins. ***B,*** A parsimonious model that partially explains the copy number and rearrangements found in the AE54SL chromosome 10. Assuming that at some point there was chromosome 10 aneuploidy (see Supplementary Figure 4), the observed DOC indicates the presence of one intact chromosome, while two or even 4 more copies formed isochromosomes. Such event(s) would create a partial monosomy of the 6 Mb region at the 5’end of chromosome 10 and a trisomy of the remaining ~8.7 Mb portion. At least one putative isochromosome has the 354bp unique loop (colored blue) and ~300 kb deletion (mR) in one arm. The further rearrangements are difficult to reconstruct, in particular the occurrence of several regions after position 6,003,515 with high depth of coverage. The high copy number region harboring integrated vector (chromosomal position 6,863,078) is unstable and undergoes constant copy number change (see Supplemental Information, Figure 4), which might result in the head-to-head duplication of the vector and adjacent upstream chromosomal region after the initial vector integration.

Figure 4 illustrates some of the CGRs in chromosome 10 of this clone. Comprehensive analysis of this region revealed vector pMGX (Figure 3) was integrated into one copy of the amplified ~220 kb fragment at chromosomal position 6,863,078. Multiple copies of this region are also present in the AE63XG genome, which is the parental cell line used for transfection with pGMX to generate the AE54SL clone, though its copy number in the AE54SL chromosome 10 is higher (approximately 10 vs 7 copies; see Supplementary Figure 2). The subsequent inverted vector duplication probably occurred upon replicative recombination within the expanding repeat with concurrent loss of the 3’ flanking chromosomal sequence. The whole vector-containing ~220 bp segment then underwent a duplication. The WGS sequencing read coverage of the pMGX vector is consistent with its four copy number presence in the AE54SL genome (Table 2).

#### AD49ZG and AD49ZH cell lines

The DNA from these cell lines was analyzed by CRISPR/Cas9-based RGEN-TdT protocols and the results were compared with the hybridization capture xGen protocol, TLA, and with WGS (data sets generated using both Illumina® and PacBio systems) (Table 2). The WGS data suggests there are two pCLD1 vector copies in the AD49ZG (Table 2) cell line, which is confirmed by qPCR (data not shown). Breakpoints within a 2.3 Kb direct repeat in the vector (Figure 5) could be redundant (pairs 3,601, 6,580, and 4,241, 7,223). The xGen protocol has identified only one host|vector junction (chr7|758) and three vector|vector junctions: 548|11,277, 3,599|11,208, and 8,348|8,330. The same host|vector junction and one out of three vector|vector junctions (3,599|11,208) were detected by the RGEN-TdT and WGS (Illumina® and PacBio) protocols. Finally, we inspected all identified breakpoints using the alignment visualization tool Ribbon^25^ and confirmed only one of the three vector|vector junctions (see Supplementary Figures 5, 6). Thus, the other two vector|vector vector fusion breakpoints in the xGen data (i.e., 548|11277, 8348|8330) appear to be PCR chimeras produced during the two rounds of library amplifications used by this protocol. Similarly, multiple false positive fusion breakpoints within the TLA data set may have caused difficulty in determining the precise host-vector junction in the AD4489ZG clone via the TLA method.

**Figure 5.**
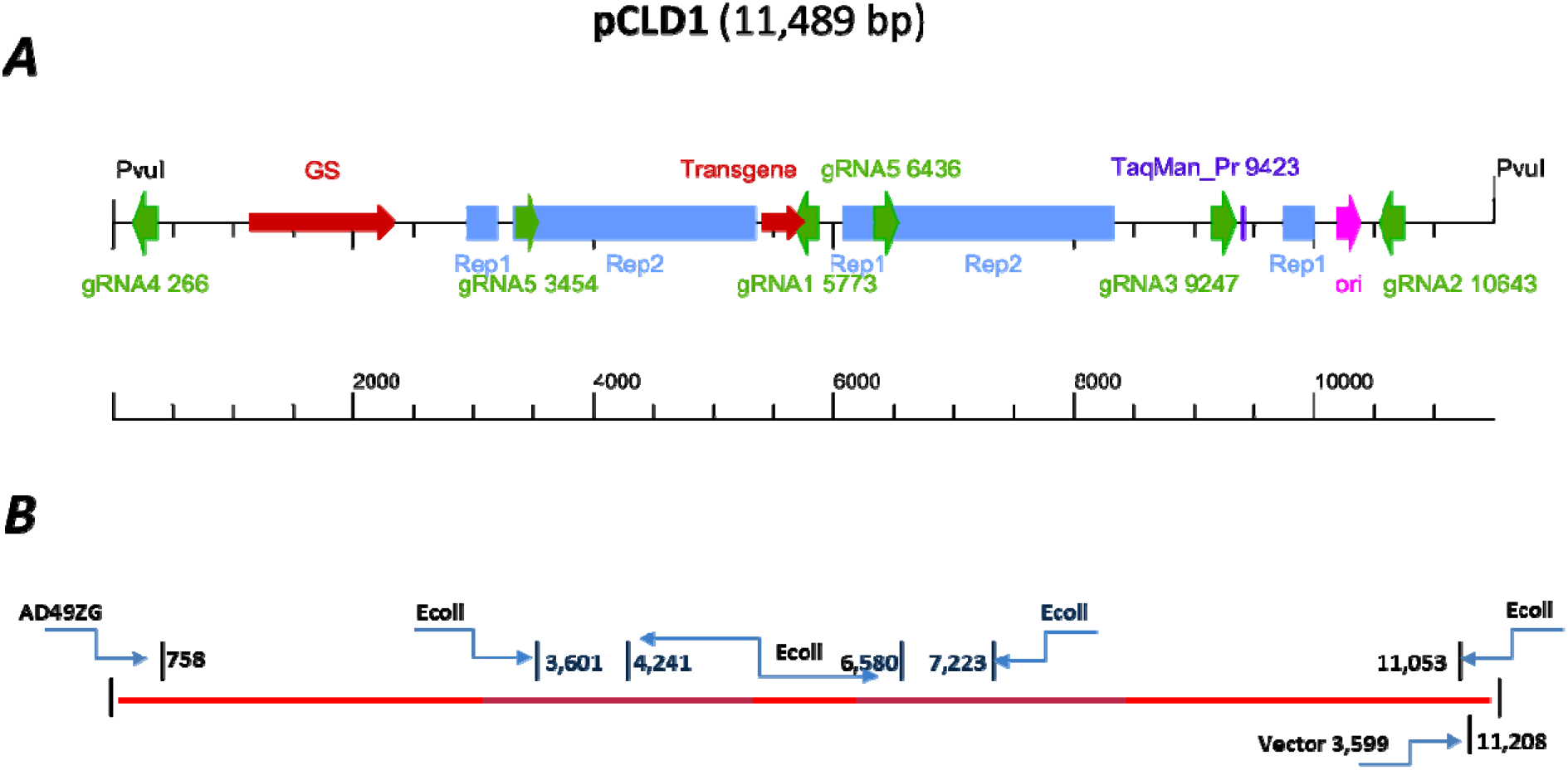
Map of the integration vector pCLD1 with breakpoints in the AD49ZG cell line. ***A,*** Different vector features are shown color-coded. Sizes of guided RNAs used in the RGEN-D protocol are not to scale. All gRNAs used in the RGEN-D protocol are shown; only gRNA1 was used for the RGEN-TdT protocol. ***B,*** Identified fusion breakpoints are shown. Blue boxes indicate direct repeat. Fusion breakpoints with *E. coli* sequences are noted (see text).

We were intrigued by the presence of the vector|*Escherichia coli* (*E. coli*) fusion breakpoints in all AD49ZG datasets. Additional analysis of the WGS data revealed about 150 Kb of an *E. coli* K-12 MG1655 genome co-integrated with the pCLD1 vector into chromosome 7 in an intricate way, resulting in the ~355 Kb integration locus (Figure 6; Supplementary Figures 7, 8). This suggests that fragments of contaminating *E.coli* DNA in plasmid preparations were co-integrated with the vector. A similar finding was recently reported during the analysis of mouse transgenic sites using the TLA method^26^.

**Figure 6.**
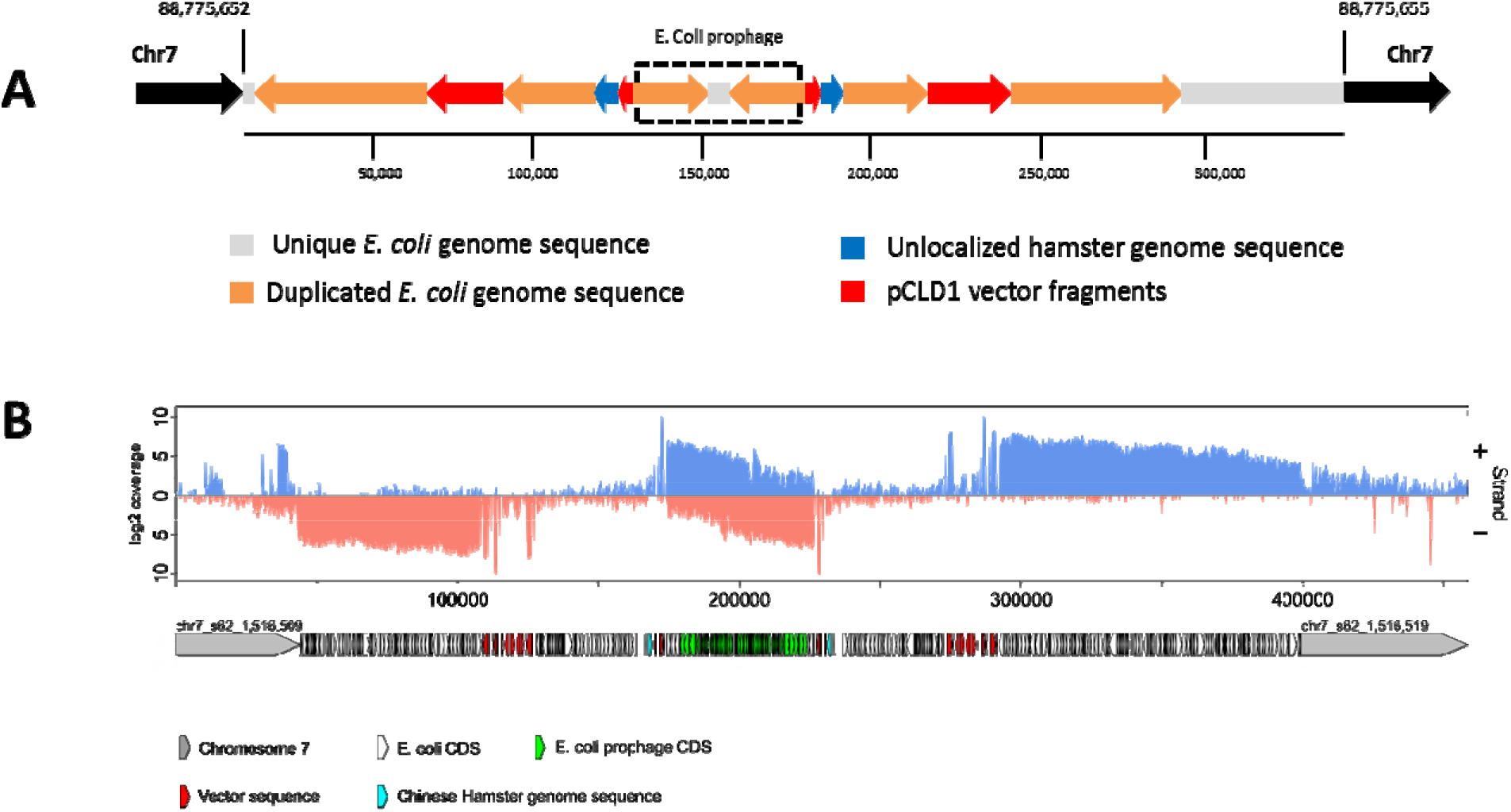
***A,*** Structure of the integration site in AD49ZG genome. The integrated sequence was assembled using Illumina® and PacBio WGS datasets (see Supplemental Information for details). ***B,*** Coverage of transcription reads aligned to the assembled AD49ZG insert with flanking chromosome 7 sequences. The insert sequence was annotated by Rapid Annotations using Subsystems Technology (RAST)^27^ (Supplementary Table 3 and Supplementary Figure 11).

It is interesting that genes encoded by the insert show high level of transcription as revealed by RNA sequencing of total RNA isolated from AD49ZG cells (see Methods). It appears that transcription of insert genes is driven by vector-encoded transgene promoter (Figure 7). However, only three reliably detected proteins out of a total of 5,358 identified by LC-MS/MS (Supplementary Table 4) are encoded by the insert, and all three are derived from the vector fragments of the insert (*C. griseus* glutamine synthetase (GS), *E. coli* β-lactamase, and an *E. coli* 55 kDa hypothetical protein; Figure 6*A*). There might be also a forth protein derived from the *E. coli* part of the insert (RNA 2’-phosphotransferase) but the spectral count of this protein is too low to be confident in its expression (Supplementary Table 4). Though the significance of the integrated and transcribed *E. coli* sequences is unclear, prior reports have shown that bacterial sequences covalently attached to expression cassettes can contribute to gene silencing^28^.

Analysis of the related AD49ZH clone (transfected with the same pCLD1 vector) revealed two host|vector junctions in chromosome 3 (84,250,935|<-499 and 486 <-|124888077) and no vector|vector fusions. It is worthy to note that data generated with the xGen protocol suggests another vector concatenation junction 8,525|8,548, which is unsupported by both the RGEN-TdT protocol and WGS, and therefore is likely a PCR artifact. TLA data also supports the same two host|vector junctions in chromosome 3 in the AD49ZH clone but also, report multiple false positive vector breakpoints (data not shown). While the two host|vector fusion breakpoints are in different scaffolds (likely both corresponding to regions of chromosome 3, data not shown), the chromosomal coverage around the breakpoints is over 30X of the average genome average, suggesting that the integration happened in a misassembled amplified chromosomal region. Since pCLD1 is present in 20 copies in AD49ZH, it is possible that a vector monomer have integrated into a single repeat unit, which expanded further (20X) after this event.

## DISCUSSION

This study explores the enrichment of long DNA fragments from mammalian genomes using the CRISPR/Cas9 system for specific targeting of genomic loci. Compared to other popular targeted techniques, like hybridization-based xGen or TLA, our RGEN protocols do not use PCR amplification and thus are essentially free from PCR-introduced chimeric junctions. Being able to enrich for long DNA fragments, the RGEN protocols give a wider range of possibilities for sequencing of complex genomic regions that cannot be studied with other methods. While we only used Illumina’s® sequencing technology to assess the RGEN methods, the isolated fragments may also serve as suitable input material for the third-generation long-range sequencing systems like Oxford Nanopore’s MinION system.

We have identified several bottlenecks in these protocols, which if eliminated, could increase the target enrichment substantially. For example, the RGEN-R protocol would benefit from optimizing the oligonucleotide duplex ligation to long genomic fragments and from improved restriction digestion of the ligated duplex. Also, the binding specificity and efficiency of biotinylated multi-kilobase DNA fragments to streptavidin beads, as well as the fragment recovery, needs to be improved in all protocols.

We have demonstrated the utility of the RGEN protocols by capturing and determining, through sequencing, the fusion breakpoints in loci of three CHO-derived cell lines that are associated with integrated vectors. The seemingly simple task of identifying vector integration boundaries in hamster genomes is not as simple as in human or mouse cell lines. This is because of a lack of a high quality assembly of the Chinese hamster or CHO genomes^29^ and because of the inherent nature of aneuploidy and continuing chromosomal rearrangement in CHO cells^14,30^. The cell lines investigated in this work illustrate the complexity of this problem and highlight the limitations of target enrichment protocols in general. We were able to determine structures of integration loci only in combination with whole genome sequencing.

Interestingly enough, in two out of three CHO-K1 cell lines studied here vector integrations occurred in unstable high copy number chromosomal regions (see Supplementary Figures XX). If this is a wide spread phenomenon, then such recombinant transgene producing clones may undergo further rearrangements involving the integration site^31^, with unpredictable phenotypic consequences. Given our findings, and the different chromosomal stability in CHO cells^16^ (Supplementary Figures 2, 4) it might be attractive for the industry to move away from making recombinant CHO cell lines by random transgenesis and adopt CRISPR/Cas9 assisted locus targeting or other precise gene targeting modalities^32^ as more robust approaches to construct stable cell lines. Identification of neutral loci on stable chromosomes appears to be attractive candidate locations for establishing high-yielding CHO cell lines.

## METHODS

### CHO-K1 Cell Lines and Clones

Plasmid pCLD1 (^~^20ug) was electroporated into ^~^5×10^6^ CHO K1 GS-/-cells. Upon recovery from transfection, cells were plated in 96 well plates at ^~^5,000 cells per well in 20% EX-CELL CD CHO Fusion Media/80% EX-CELL CHO Cloning Media. Stable IgG expressing pools were established via selection in glutamine deficient media and IgG secretion was evaluated via bio-layer interferometry (BLI). The highest expressing pools were used for cloning via limiting dilution in 20% conditioned media/80% EX-CELL CHO Cloning Media. Clonality was determined based upon visual inspection five days post-plating and IgG secretion of all clones was monitored via BLI. The top expressing clones were scaled up, banked and tested for stable IgG secretion in a 70-generation stability assay.

### Primers and sgRNAs

DNA sequence information from the genome assembly CHOK1GS_HDv1^33^ (GCA_900186095.1) of the Chinese hamster ovary cell line CHOK1 was used for designing guide RNAs. The following web resources were used for selecting primers and crRNA sequences: http://chopchop.cbu.uib.no (a webtool for crRNA design^34^), NCBI’s Primer-BLAST https://www.ncbi.nlm.nih.gov/tools/primer-blast/, and ThermoFisher’s Multiple Primer Analyser. All oligonucleotides (both modified and unmodified ones) were synthesized by IDT, TaqMan probes were ordered from ThermoFisher. Sequences of primers and crRNA target sites are provided in Supplementary Table 1. Single-guide RNAs were prepared by IDT’s method; custom synthesized 36-mer crRNAs were annealed to universal 67-mer tracrRNA. In the RGEN-D protocol tracrRNA contained biotin at the 3’-end.

### Comparative C_T_ (threshold cycle) method of real-time PCR

In this method^18^, which is typically used for gene expression profiling, the real-time PCR data is presented relative to another gene which is often referred to as an internal control. Accordingly, PCR amplicons were designed for the vectors that either span the RGEN cut site inside the vector sequence or are positioned in close proximity to the cut site to serve as an internal control. Upon cleavage of gDNA with RGEN(s), the amount of PCR amplicon spanning the RGEN cut sites diminishes depending on the efficiency of RGEN cleavage, while upstream and/or downstream internal control amplicons stay unaffected, thus allowing calculation of percentage of cleavage. An example of this method for evaluation of the RGEN-1 cleavage efficiency is provided in the Supplementary Figure 12.

### Comparative ∆∆C_T_ TaqMan Assay

This assay was designed for estimating the degree of depletion of non-target sequence *versus* target sequence as a result of the capture. Based on this assay, a decision is made about NGS library construction. Two TaqMan probes are used in the assay, one designed for a target region, which is the common region in the transgene sequences in vectors pCLD1 and pMGX (ThermoFisher’s TaqMan assay Hs00378230_g1), while the other is chosen outside of the target region (*Cgr* Spag5 gene region; ThermoFisher’s Taqman assay Cg04462858_g1) and serves as an internal control. TaqMan probes in the assays are labeled with different fluorescent dyes (FAM™ and VIC™) and can be conveniently used on the same DNA template in one tube. This greatly decreases all experimental errors associated with pipetting and using separate tubes. The following equation is used to compare the target sequence enrichment in two different samples (untreated and treated DNA samples):

Fold change = 2^-∆∆Ct^ = [(C_T_ target sequence amplicon – C_T_ internal control amplicon)treated DNA – (C_T_ target sequence amplicon – C_T_ internal control amplicon)untreated DNA)].

### DNA Preparation for NGS in RGEN-A Protocol

Genomic DNAs were isolated using QIAGEN’s EZ1 and DNeasy Blood & Tissue kit. To digest target sequences in the genome, Cas9 protein (2–10 μg) (New England Biolabs (NEB)), which had been preincubated with sgRNA1 at room temperature for 15 minutes to make RGEN-1, was mixed with genomic DNA (10-20 μg) in a Cas9 reaction buffer with added 1 M betaine and 50 μg/ml bovine serum albumin (BSA) and incubated at 37°C for 8 hours. Digested genomic DNA was purified again after incubation with RNase A (50 μg/mL) for 1 hour at 37°C followed by incubation with Proteinase K (50 μg/mL) for 1 hour at 50°C. A single ‘A’ nucleotide was added to the 3’ ends of the blunt fragments using NEB’s A-tailing protocol with Klenow Fragment. A corresponding single ‘T’ nucleotide on the 3’ end of the adapter provides a complementary overhang for ligating the adapter to the fragment. The duplex adaptor (1 pM) with a biotinylated 5’-end and containing a site for rare cutting restriction endonuclease AsiSI was ligated to A-tailed DNA ends using the Quick Ligation Kit (NEB). Double stranded biotinylated DNA molecules were then immobilized to streptavidin magnetic beads using Dynabeads kilobaseBINDER kit (ThermoFisher) according to the manufacturer’s recommendations, and then subsequently released by digestion of the ligated adaptor with AsiSI. The enrichment of the targeted locus was estimated by comparative ∆∆C_T_ TaqMan assay.

### DNA Preparation for NGS in RGEN-TdT Protocol

Isolation and digestion of genomic DNA with RGEN-1 was performed as described in the RGEN-A protocol. In the initial variant of this protocol, to demonstrate the specific capture and sensitivity of DNA strand breaks in vitro, RGEN-1 treated DNA was directly labeled by the terminal deoxynucleotidyl transferase-mediated dUTP-biotin end-labeling (TUNEL) using a recombinant terminal transferase (TdT) and according to the standard tailing protocol from the supplier (NEB). Briefly, 15 μg of digested gDNA was incubated in TUNEL reaction mix containing 5 mM CoCl_2_, 0.20 mM of biotin-16-dUTP, and 1600 U of TdT at 37°C for 1 hour in a final volume of 70 μL. The end-labeling reaction was arrested by heating the reaction mix at 70°C for 10 min. To remove TdT, Zymo Research’s DNA Clean & Concentrator-25 columns were used. It was noticed that this step may result in significant loss during purification. As an alternative LiCl/ethanol precipitation can be used. Labeled DNA molecules were immobilized to streptavidin magnetic beads using Dynabeads kilobaseBINDER kit (ThermoFisher) and then subsequently released according to the protocol from the supplier. After purification with QIAGEN’s MinElute PCR purification kit, DNA was double-stranded/amplified with Illustra GenomiPHI kit (GE Life Sciences) and the enrichment was evaluated using comparative ∆∆C_T_ TaqMan assay. The enrichment of the targeted locus using the improved RGEN-TdT method was estimated by comparative ∆∆C_T_ TaqMan assay.

### DNA Preparation for NGS in RGEN-D Protocol

Instead of non-labeled tracrRNA, tracrRNA biotinylated at the 3’ end was used. To make individual biotinylated sgRNAs, equal volumes of tracrRNA-3’-biotin and each of crRNAs were incubated at 95°C for 2 minutes and cooled down to room temperature. dCas9 (NEB, EnGen Epy dCas9) protein (2–10 μg), was then incubated with equimolar mixture of sgRNA1-5 (vector pCLD1) or sgRNA6-10 (vector pGMX) (Figures 5, 6 and Supplementary Table 1) at room temperature for 15 minutes to make the corresponding RGENs. Sonicated genomic DNA (10-20 μg, 5 kb average size) was mixed at room temperature for 20 minutes with RGENs conjugated to 50 μL of Dynabeads-Streptavidin M280 beads. RGEN-Dynabeads conjugation was performed in Cas9 reaction buffer with added 1 M betaine and 50 μg/mL BSA, at room temperature for 15 minutes. RGEN-Dynabeads with bound DNA were washed four times with 0.3 mL of modified low salt buffer in the presence of 0.05% Triton. The captured RGEN-DNA complexes were treated with RNase A and Proteinase K. DNA was purified using ChIP DNA Clean & Concentrator (Zymo Research). Alternatively, bound DNA was released from the beads by treatment with 95% Formamide at 65°C for 5 minutes followed by DNA purification using microspin G-50 colums (GE Lifesciences). Released DNA was amplified using the Illustra GenomiPhi kit (GE Lifesciences) before proceeding with NGS library preparations.

### Construction of Genomic Libraries for NGS and Sequencing

Captured DNA (AE54SL, AD49ZG, and AD49ZH clones) and genomic DNA (clone AE63XG) fragmentation was performed using a Covaris S220 system and appropriate consumables and protocols. Typically, DNA was fragmented to 500 bp. Fragmented genomic DNA was ligated with adaptors using Illumina’s TruSeq Nano DNA Library Prep kit or the Nextera XT kit in case of xGen protocol (IDT). The concentration of the libraries was determined using the KAPA Library Quantification Kits for Illumina platforms (Kapa Biosystems). Libraries were sequenced using an Illumina® MiSeq sequencer in a paired-end mode with 301 (in some cases with 76 and 251) cycles per read. For Nextera XT libraries, the DNA was enzymatically “tagmented” in lieu of fragmentation. The PCR duplication rate in xGen protocols was calculated using a computational method of Bansal^24^ to estimate the average PCR duplication rate that accounts for natural read duplicates by leveraging heterozygous variants in a genome.

PCR-free libraries of AD49ZG and AD49ZH genomic DNAs were prepared and sequenced by Macrogen Inc. One library per sample was constructed using Illumina’s TruSeq PCR Free Library Preparation kit with insert size of 550bp. Sequencing was performed using the Illumina HiSeq2500 system with 250bp paired-end sequencing method. Each library was sequenced over one entire flowcell, which contains two lanes. Each flowcell produced over 700M reads and resulted in ~70X coverage for both DNA samples.

For PacBio SMRT sequencing of the AD49ZG genomic DNA, SMRTbell DNA template libraries (insert size of ~20 Kb) were prepared according to the manufacturer’s specification followed by size selection using Sage Science’s BluePippin instrument to remove short molecules. SMRT sequencing (10 SMRT cells) was performed on the Pacific Biosciences Sequel System by Macrogen Inc.

### Reconstruction of CHOK1 Chromosomes

For the characterization of integration loci we used *Cricetulus griseus* GenBank assembly GCA_000448345.1 (Cgr1.0) and the most recent draft genome assembly CHOK1GS_HDv1 (GenBank assembly accession GCA_900186095.1) of the Chinese hamster ovary cell line CHOK1^33^. The last assembly contains the fewest number of scaffolds compared to previous CHO cell lines assemblies^35–38^; in some cases scaffolds approach the size of individual chromosomes. Brinkrolf et al^35^ isolated, by flow cytometric cell sorting, *Cricetulus griseus* (*C. griseus*) chromosomes and sequenced them individually. This has made possible the assignment of scaffolds to most of *C. griseus* chromosomes (Cgr1.0 assembly^35^). Using their data as well as data of Wlaschin and Hu^39^, corresponding scaffolds from the CHOK1GS_HDv1 assembly were concatenated to make chromosomes 1-8, and X. Because chromosome 1 exceeds 2^29^ bp, it was split into two equal parts (referred to as chr0 and chr1) for compatibility with different bioinformatics tools used for the analysis. Remaining scaffolds belonging to chromosomes 9, 10, and Y were not assigned to individual chromosomes in the original work because chromosomes 9 and 10 could only be separated as a pool and chromosome Y was not sorted at all during sequencing^35^. We found, however, a way to assign scaffolds between chromosomes 9 and 10 and properly orient them within respective chromosomes *in silico*, using corresponding CHOK1GS_HDv1 scaffolds, a good synteny of these scaffolds with an internal region of the mouse chromosome 15, and our data on chromosomes coverage (Supplementary Figures 1). Scaffolds belonging to chromosome 7 were also oriented based on the synteny with mouse chromosomes 11, 12, 16, and 17 (Supplementary Figure 2). All other unplaced scaffolds were combined into pseudochromosome Y.

### Processing DNA Sequencing Data

Illumina NGS data analysis followed an established approach that included (i) raw data pre-processing, (ii) mapping of reads to reference genome, and (iii) extracting mapping information and sequence context of vector integration site(s). A customized reference genome for each CHO cell clone represented a fasta sequence file of reconstructed CHO-K1 chromosomes with a corresponding vector sequence and *E. coli* genome (RefSeq NZ_CP017100.1) as a pseudochromosomes. We found that external data pre-processing (*i.e*., adapter and quality trimming) can be safely skipped as adapter trimming can be done by the instrument software (*e.g*., MySeq Reporter) during raw data processing, and quality trimming was generally not required. Thus, in most cases machine generated fastq files were directly used for mapping to reference genomes. For mapping Illumina sequencing reads and for extracting and visualizing sequence context of vector integration sites, Burrows-Wheeler Aligner (bwa mem)^40^, samtools^41^, bbmap package^42^, and IGV^43^ or Consed^44^ programs were used. An example of a minimal analysis session is provided in the Supplemental Information. We also used Ribbon^25^ and SplitThreader^45^ for validation of structural variations.

For building reference depth of coverage (DOC) maps of *C. griseus* chromosomes, we downloaded from NCBI SRA archive 336 M reads (BioProject PRJNA167053) and mapped them to our customized reference genome (without vector and *E. coli* pseudochromosomes).

PacBio sequencing reads were aligned with NGM-LR^46^ and SVs were called with Sniffles^46^, which is a structural variation caller using third generation sequencing (PacBio or Oxford Nanopore). It detects all types of SVs (10bp+) using evidence from split-read alignments, high-mismatch regions, and coverage analysis.

### Assembly of Integrated Sequence in AD49ZG genome

Illumina HiSeq2500 251-bp paired reads (total 713,405,402 reads; Supplementary Table 2) were mapped with bwa mem to a combined reference sequence containing the CHOK1GS_HDv1 assembly, *E. coli* K-12 MG1655 genome (Genbank accession number NZ_CP017100.1), and the pCLD1 vector sequence (Supplementary Figure XX). Similarly, starting with 10 SMRT cells, 5,210,389 uncorrected continuous reads longer than 1000 bases were obtained, merged together into a single dataset, and mapped with NGM-LR^46^ to the combined reference sequence. Using samtools^41^ to identify read names from an Illumina bam file mapped to *E. coli* and pCLD1 sequences, a unique list of corresponding read pairs was created and then used by BBTools^47^ to extract paired read sequences from the HiSeq2500 dataset. Pacbio subreads mapped to *E. coli* and pCLD1 sequences were extracted in fasta format from a corresponding bam file in a similar way. The total 44,850 vector-and E. coli-mapped HiSeq2500 reads were then assembled by spades^48^ (v. 3.11.1), while 326 Pacbio subreads were assembled separately by canu^49^; spades was also used for a hybrid assembly of the pCLD1-*E. coli* insert using both HiSeq2500 reads and Pacbio subreads. The final insert sequence was produced in Consed^44^ after manual evaluation of three different assemblies (Supplementary Figures 9, 10).

### AD49ZG Transcriptome Sequencing and Analysis

Total RNA was extracted from 10^6^ AD49ZG cells using the RNeasy Mini kit (QIAGEN) following the manufacturer’s protocol. The purity and yield of total RNA was analyzed using the Quant-It™ RNA Assay Kit (Life Technologies) and SpectraMax M2 plate reader (Molecular Devices). The RNA integrity was confirmed by the Agilent 2100 Bioanalyzer system (Agilent Technologies). Two samples of 1 μg each with an RNA integrity number value greater than 8 were used for preparation of sequencing libraries using TruSeq™ Stranded Total RNA Sample Preparation Kit (Illumina®) and protocol (fr-firststrand libraries). The library concentration was determined with the KAPA Library Quantification kit for Illumina platforms (Kapa Biosystems). RNA sequencing was performed using a paired-end strategy (each end with 300 bases) on the Illumina® MiSeq instrument.

Adaptor trimmed fastq files from two MiSeq runs were combined (total 117 M reads) and second reads (read2) were mapped to the customized AD49ZG genome containing CHO-K1 chromosomes and the ^~^320 kb assembled integration sequence. Reads were aligned to the reference with Burrow-Wheeler Aligner (BWA)^40^.

### AD49ZG Total Protein Profiling with LC-MS/MS

#### Sample Preparation

The cell pellet (10^7^ cells) was suspended in 400 μL of modified RIPA buffer (2% SDS, 150mM NaCl, 50mM Tris.HCl pH8, 1X Roche Complete protease inhibitor) and sonicated using a sonic probe (QSonica) with the following settings: Amplitude 50%, Pulse 10 x 1s. 1 on 1 off. The protein concentration of the lysate was determined by Qubit fluorometry (Invitrogen). 20μg of protein was processed by SDS-PAGE using a 4-12% Bis-Tris NuPAGE mini-gel (Invitrogen) with the MOPS buffer system. The gel lane was excised into forty equally sized bands and in-gel digestion with trypsin was performed on each using a ProGest robot (Digilab) with the protocol outlined below.

- Washed with 25mM ammonium bicarbonate followed by acetonitrile
- Reduced with 10mM dithiothreitol at 60°C followed by alkylation with 50mM iodoacetamide at room temperature
- Digested with trypsin (Promega) at 37°C for 4 hours
- Quenched with formic acid and the supernatant was analyzed directly without further processing.

#### Mass Spectrometry

Half of each digested sample was analyzed by nano LC-MS/MS with a Waters NanoAcquity HPLC system interfaced to a ThermoFisher Q Exactive mass spectrometer. Peptides were loaded on a trapping column and eluted over a 75μm analytical column at 350nL/min; both columns were packed with Luna C18 resin (Phenomenex). The mass spectrometer was operated in data-dependent mode, with the Orbitrap operating at 70,000 FWHM and 17,500 FWHM for MS and MS/MS respectively. The fifteen most abundant ions were selected for MS/MS.

#### Data Processing

Data were searched using a local copy of Mascot with the following parameters:

- Enzyme: Trypsin/P
- Database: UniProt Chinese hamster reference proteome appended with the target sequences encoded by 315 kb insert
- Fixed modification: Carbamidomethyl (C)
- Variable modifications: Oxidation (M), Acetyl (N-term), Pyro-Glu (N-term Q), Deamidation (N,Q)
- Mass values: Monoisotopic
- Peptide Mass Tolerance: 10 ppm
- Fragment Mass Tolerance: 0.02Da
- Max Missed Cleavages: 2

Mascot DAT files were parsed into Scaffold (Proteome software) for validation, filtering and to create a non-redundant list per sample. Data were filtered using at 1% protein and peptide FDR and requiring at least two unique peptides per protein.

## ACKNOWLEDGEMENTS

We thank Jaya Onuska for her help with DNA samples preparation, Sindy John and members of her group for bioinformatics assistance. We also thank Sergei Kozyavkin for his thoughtful comments on the results and David Martin for his excellent IT support and help with Google cloud computing.

## AUTHOR CONTRIBUTIONS

Author contributions: A. S., C.C., A.C., and D.O. designed research; T.B., K.A., and D.R. contributed new CHOK1 cell lines; L.V. constructed transcriptome libraries; A. S., C.C., and Y.T. analyzed data; and A.S., C.C. wrote the paper.

## DATA AVAILABILITY

Complete sequencing data have been deposited in the Sequence Read Archive (SRA) under the accession number SRP151278.

